# A shotgun approach to identify mechanical nociception genes

**DOI:** 10.1101/077644

**Authors:** Melissa G. Christianson, Stephanie E. Mauthner, W. Daniel Tracey

## Abstract

The molecular mechanisms of sensing noxious mechanical force by nociceptive sensory neurons remain poorly understood. Traditional methods for probing mechanical nociception behavioral responses are labor intensive and involve the testing of one animal at a time. This time consuming process of behavioral testing has largely precluded large scale analyses. Indeed, large scale genetic screens that have been performed thus far have been largely restricted to the investigation of ion channel genes [1]. Here we describe a new behavioral assay for mechanical nociception in which tens of animals can be stimulated simultaneously. In this assay, third instar larvae of the genetically tractable organism *Drosophila melanogaster* are mechanically stimulated with tungsten particles that are fired from a gun. We have used the new assay to carry out a genetic screen in which we investigated the function of 231 nociceptor enriched genes with tissue-specific RNA interference. Targeting of 21 genes resulted in mechanically insensitive phenotypes and targeting of a single gene resulted in a hypersensitive mechanical nociception phenotype. Six of the identified genes were previously uncharacterized and these were named after famed Roman gladiators (*Spartacus (CG14186*), *Commodus (CG1311), Flamma (CG10914), Crixus(CG6685), Spiculus (CG10932),* and *Verus (CG31324)*).

## Introduction

*Drosophila* larvae offer an excellent system in which to study molecular pathways responsible for nociception. The neurons important for transducing nociceptive stimuli in *Drosophila* larvae–called Class IV multidendritic (md) neurons–are known [2], and they resemble vertebrate nociceptors in important functional ways. Importantly, both use transient receptor potential channel (TRP) family ion channels to generate neurobehavioral responses [3–6] and both become sensitized following injury [7, 8]. Several nociception pathways have found to be conserved between *Drosophila* and mammals [3, 8–12]. The existence of powerful genetic tools in concert with the rapid *Drosophila* life cycle allows the study of *Drosophila* nociception *in vivo* in ways that would be impractical or impossible in other model systems.

Third-instar larvae respond to noxious mechanical or thermal stimulus with a stereotyped response involving corkscrew rotation[3] about the long-body axis that is followed by a period of rapid crawling[13]. Assays of thermal and mechanical nociception have been developed to investigate these behavioral responses[3]. However, these assays are laborious and time-intensive, requiring individual stimulation of single animals and assessment of subsequent behavior. High-throughput methods for avoidance of high temperature in larval and adult *Drosophila* have been reported [12, 13], but no method specifically probing mechanical nociception exists.

As a result, even though the polymodal Class IV md neurons are required for sensation of all or most types of noxious stimuli [2, 5, 11], the extent of overlap between the pathways for mechanical and thermal nociception is poorly understood. *painless* and *dTRPA1,* which both encode TRP channels, are functionally important for both thermal and mechanical nociception[3, 5, 10, 14]. The molecular pathway for mechanical nociception requires *pickpocket (ppk)* [11, 15] and *balboa(bba)/ppk26* which both encode ion channels of the DEG/ENaC family[16–18] and the *Drosophila piezo* gene[15]. The PPK/BBA heteromeric channel is required for sensing noxious mechanical stimuli, but it is not required for thermal nociception or for optogenetically triggered nociception behaviors[11] [18]. Surprisingly, *ppk-RNAi* actually causes hypersensitive thermal nociception responses and at the same time cause insensitive mechanical nociception responses which indicates that single genes can affect thermal and mechanical nociception in opposite ways [19]. Given this evidence that genes show specific effects on mechanical nociception pathways and thermal nociception pathways–screens relying solely on tests for thermal responses are likely to miss many of the important players in mechanical pathways and vice versa. Whereas thorough studies of genetic pathways essential for thermal nociception have been made in *Drosophila* larvae [3, 19] and adults[12], a counterpart investigating the response to noxious mechanical stimuli has yet to be performed.

Therefore, we sought in this work to identify genetic pathways mediating *Drosophila* mechanical nociception responses. To do so, we have developed a new and relatively high-throughput method. Using this method we screened through a set of nociceptor-enriched genes that were identified by laser capture microdissection and microarray analyses [18, 19]. The development of the new paradigm and the results of the screen are reported here.

## Results

Because currently available methods assaying mechanical nociception behaviors in *Drosophila* larvae involve laborious methods that test individual animals one at a time [20], we developed a method that instantaneously delivers a noxious mechanical stimulus to a population of animals. To do so, we placed wandering third instar larvae in a behavioral arena and ballistically stimulated (shot) them with 12-µm tungsten particles emitted from a gene gun (Figure 1A).

**Figure 1.**
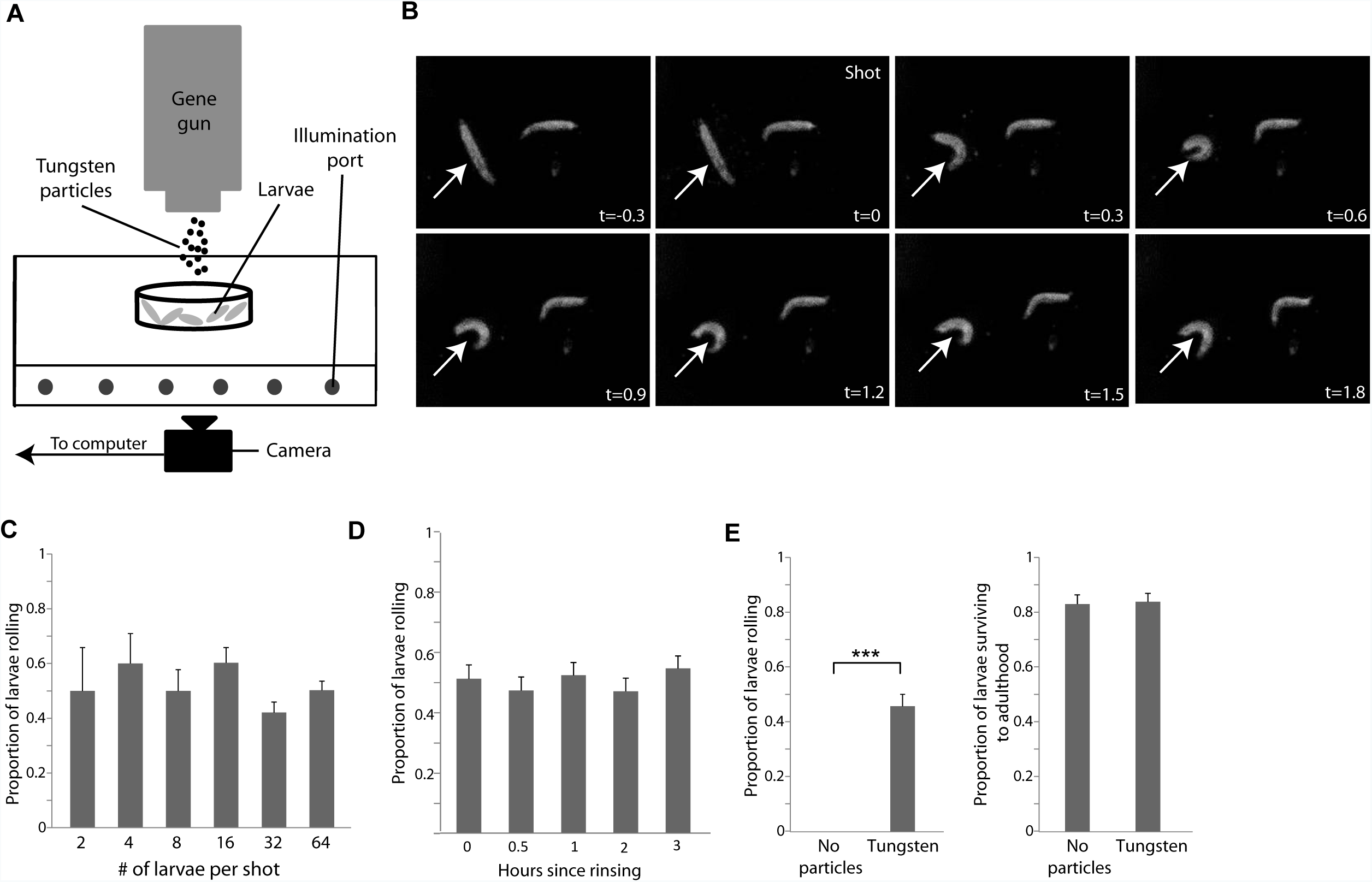
Tungsten particles fired from a gene gun cause nocifensive escape locomotion in *Drosophila* larvae. **(A)** Schematic showing the behavioral apparatus used for the gunshot assay with wandering 3^rd^ instar *Drosophila* larvae. **(B)** Larval images extracted from digital video recordings showing the typical larval nocifensive escape locomotion response (*arrow*) to the gunshot. **(C)** Gunshot assay with increasing numbers of w1118 larvae stimulated per shot demonstrating no change in larval responsiveness with test population density. **(D)** Gunshot assay with w1118 larvae stimulated after incubations for increasing durations of time prior to stimulation show no change in larval responsiveness over time. **(E)** Bar plots demonstrating that w1118 larvae shot with the gene gun (*left*) still develop to adulthood at a frequency similar to that of their non-stimulated counterparts (*right*). (t-test, error bars depict standard error of the proportion, *** denotes p<0.001).

Gunshot stimulated larvae exhibited a characteristic escape response of rotation of the larval body around the long body axis (rolling; Figure 1B, Supplemental Video 1), a response that was indistinguishable from the larval nocifensive escape locomotion seen with other noxious stimuli [3]. The timing of the stimulus in this assay could be precisely observed because the tungsten projectiles produced a pattern of tiny bubbles as they cavitated through the aqueous medium surrounding the larvae (Supplemental Video 1). This allowed us to observe that animals quickly initiated nociceptive responses (a short latency often within a video single frame (30 ms)) after tungsten particles reached the surface, that persisted for varying durations of time. Larvae that did not roll in response to the stimulus often engaged in other mechanosensory-related behaviors, such as turning, “scrunching”, or pausing.

In pilot studies designed to identify the most efficient settings for inducing nociception behavior with the newly developed paradigm, we fired the gun at larvae using tungsten particles of different sizes (0.5-12 µm in diameter). Small particles (<2 µm) stimulated rolling behavior less readily than large particles (12 µm) and required either a shorter gun-to-larva distance, which consequently reduced the maximum size of the behavioral arena covered by a single shot, or greater emission pressure to achieve similar response frequencies (data not shown). Thus, the 12 µm particles were chosen for use in the gunshots as this particle size efficiently stimulated rolling when fired from a distance that covered a relatively wide behavioral arena and would conceivably allow us to maximize the number of larvae we could test at once in our high-throughput assay.

To determine if the gunshot stimulus would permit simultaneous testing of many larvae, as would be necessary in a high-throughput behavioral screen, we examined behavioral responses of groups of larvae ranging from 2 to 64 in size. We found no change in individual larval responsiveness when increasing the population of larvae during testing (Figure 1C), indicating that larva-to-larva contact and possible crowding-induced mechanical stimulation are not important factors affecting the likelihood of an individual animal to roll.

High-throughput screening could require testing of several hundred genotypes of larvae in different arenas in a single testing session. Because the time necessary to assemble larvae for such screening (i.e., to rinse larvae from each genotype from food vials into separate behavioral arenas with the appropriate amount of liquid) increases linearly with the size of the screen (see Methods), we asked whether incubating larvae in behavioral arenas for varying durations of time prior to shooting would alter their propensity to roll. We found that maintaining larvae in behavioral arenas (20 mm Petri dishes) for up to 3 h prior to gunshot stimulation had no effect on larval rolling frequency (Figure 1D). We also did not observe obvious changes in larval activity or locomotion over this period.

Finally, we sought to determine the whether the gunshot stimulus was an overly severe stimulus that would impact larval survival. After engaging in nociception behavior, the stimulated larvae resumed locomotion that was not noticeably impaired and showed no notable changes in pupation time compared to mock-treated controls, suggesting that the stimulus was relatively mild but effective. We occasionally observed the appearance of melanotic spots or cuticle autofluorescence in a stimulated larva several hours post tungsten particle exposure. These spots, which are indicative of localized tissue damage and a melanization pathway [21–24], are similar to those seen after cuticle penetration by the ovipositor of parasitoid wasps, a natural noxious stimulus[14]. Finally, we found that the gunshot did not cause catastrophic larval damage leading to death, as an approximately equal proportion of stimulated larvae and mock-treated larvae completed development to eclose as adult flies (Figure 1F). Together, these data indicate that ballistic stimulation is a non-lethal stimulus of suitable intensity for high-throughput behavioral screening.

We next asked what feature of the ballistic stimulus was salient for inducing nociceptive behavior. Given that larvae did not roll when shot with a tungsten-void Helium puff (Figure 1F), we hypothesized that tungsten particles striking the larval cuticle were a critical element. Therefore, we tested whether the tungsten particle speed/acceleration (via gene gun emission pressure) and density had an effect on larval rolling rates.

The particle number density and emission pressures that were used in a reverse genetic sceen (as outlined below) resulted in approximately 30 particles/mm^2^ struck an agarose filled arena in its center (Figure 2A). We estimate this was reduced to approximately 20 particles/mm^2^ at the edge arenas used for behavioral analyses (which were intentionally smaller (20mm) than those used for our estimates of particle density (35mm))(Figure 2B). Thus, a third-instar larva which has body dimensions of about 3.5 mm in length and 1 mm in width would be struck by many particles, regardless of its location within our behavioral arenas. Consistent with this, stimulated larvae frequently had particles embedded in their cuticle (Figure 2C), with particles penetrating near the dendrites of the Class IV md neurons (Figure 2D).

**Figure 2.**
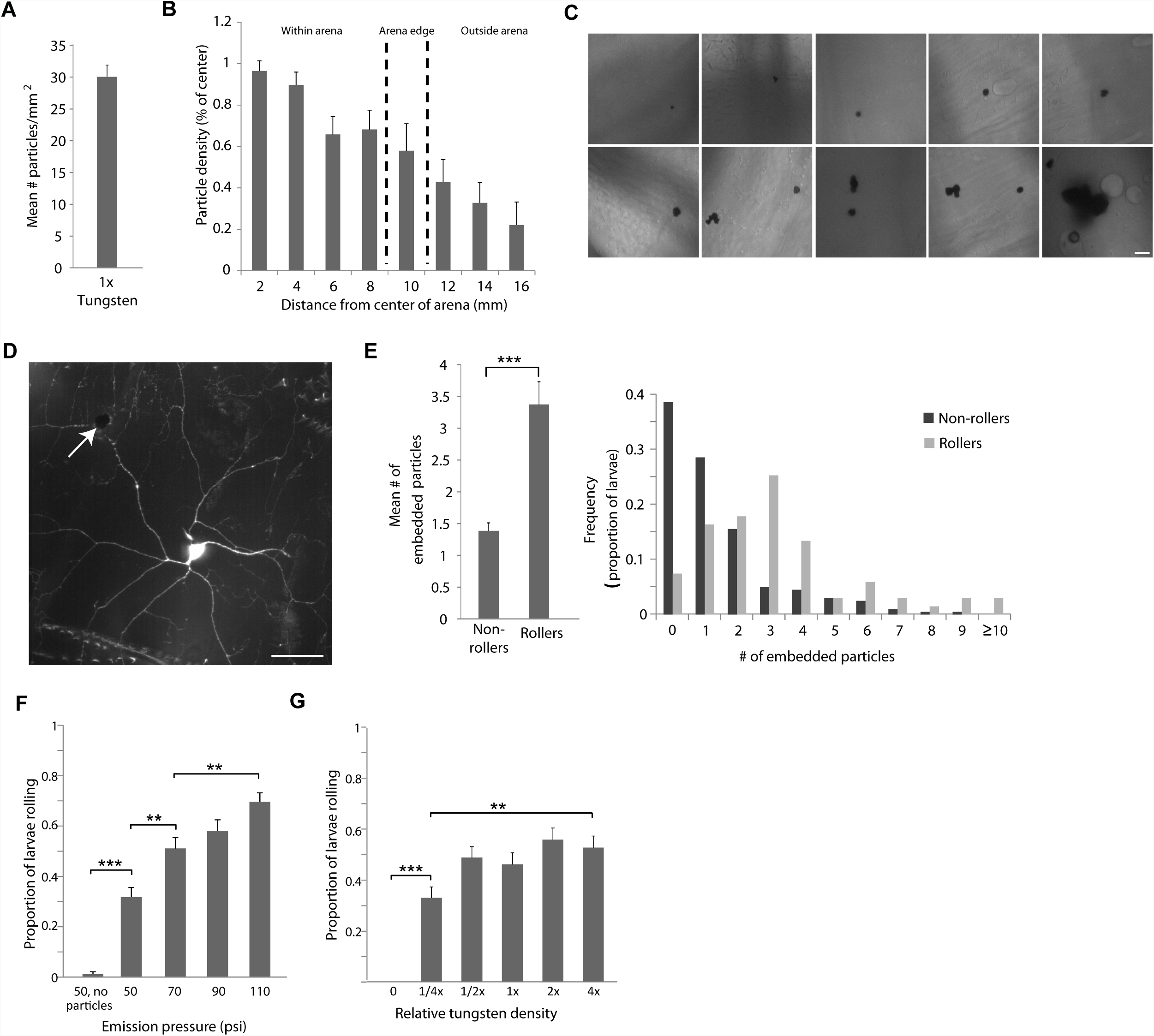
Larval rolling following the gunshot is due to tungsten particle stimulation. **(A)** Quantification of the mean number of tungsten particles in a 1-mm^2^ area at the center of the testing arena. Shot density was 1x. **(B)** Bar graph quantification demonstrating that tungsten particles are spread evenly throughout the central region of an empty testing arena. Shot density was 1/2x. **(C)** Representative photomicrographs showing particles embedded into the larval cuticle. Shot density was 1/4x. Scale bar, 20 µm. **(D)** Photomicrograph showing a particle (*arrow*) embedded near the dendrites of a mCD8::GFP-expressing Class IV nociceptor neuron in the larval body wall. Larval genotype was *ppk-GAL4 UAS-mCD8::GFP*. Shot density was 1x. Scale bar, 20 µm. **(E)** Bar graphs showing the average (*left*) and distribution (*right*) of tungsten particles penetrating into larval cuticles of *ppk-GAL4 UAS-mCD8::GFP* larvae following a gunshot stimulation at 1/4x particle density. Note that larvae rolling in response to the gunshot typically have more embedded particles than their non-rolling counterparts. **(F)** Gunshot assay using a series of increasing tungsten particle delivery pressures in *w*^*1118*^ larvae demonstrating increased larval responsiveness to stronger stimuli. **(G)** Gunshot assay using a series of increasing tungsten particle densities in *w*^*1118*^ larvae demonstrating increased larval responsiveness to stronger stimuli. (t-test, error bars depict standard error of the proportion, ** and *** denote p<0.01 and p<0.001 respectively).

The appearance of melanotic spots in the gene gun stimulated larvae suggested that at least some of the particles were capable of penetrating the cuticle and epidermis of the animals. We observed larvae with confocal microscopy following shooting in an attempt to count the number of particles embedded in the cuticle. However, using the preferred conditions of our behavioral assay for screening purposes (see below (1X tungsten and an emission pressure of 70 psi)), it was technically challenging to know unequivocally if particles attached to larvae were doing so because they had struck the larvae during the shot. This difficulty arose because following being shot the larvae were surrounded by many particles that were floating in the medium, and so we could not discern whether the particles were merely passively sticking to the larvae or if they actually struck the larvae during the shot.

Thus to overcome these uncertainties, we examined larvae that had been shot with tungsten at a very low density (1/4X tungsten) in an attempt to estimate the minimum number of embedded particles that would trigger rolling behavior. Larvae were shot with the low-density tungsten, their behavioral responses were recorded (ie rolling or non-rolling), and they were immediately observed at high magnification and the number of cuticle-embedded particles was recorded. As expected, larvae rolling in response to the gunshot typically had more particles embedded in their cuticles (Figure 2E, 3.4 ±0.4 s.e.m.) than did their non-rolling, gunshot-stimulated counterparts (Figure 2E, 1.4±0.12 s.e.m). This finding, in conjunction with the total absence of rolling in larvae stimulated with a tungsten-void air puff (Figure 1F), indicated that the tungsten striking the larval cuticle, and possibly embedding within it, was a critical factor in the triggering rolling by the gene gun stimulus.

Furthermore, at higher gene gun emission pressures, larval rolling frequency was higher than with lower emission pressure (Figure 2F). This suggests that the force-pressure relationship that is applied to the larvae when the tungsten particles struck them was an important factor in triggering rolling. Directly increasing the concentration of tungsten particles loaded into the apparatus (Figure 2G) also increased larval responsiveness. The relationship between tungsten particle number density and emission pressure with larval responsiveness highlights a convenient feature of this method as a screening tool: stimulus severity may be adjusted with relative ease.

Next, we asked whether behavioral responses to the gunshot relied on sensory neurons previously identified as key for mechanical nociception [2]. To do this, we used GAL4-UAS-mediated silencing of various md neuron classes either with the tetanus toxin light chain (to inhibit neurotransmitter release) or with RNAi against *para* (to prevent expression of a voltage gated sodium channel subunit necessary for action potential propagation). Consistent with findings from other nociception methods [2, 11], silencing Class IV neurons using either silencing method virtually eliminated gunshot triggered rolling (Figure 3A-B). Even more profound inhibition was seen with silencing all classes of md neurons (I-IV). That silencing of all md neurons shows a more pronounced effect on this nociception behavior is consistent with previous reports indicating several classes of md neuron could impair mechanical nociception to various degrees[2]. This differs from what is seen for thermal nociception in which the Class IV neurons are the only type of sensory neuron that is known to be required.

**Figure 3.**
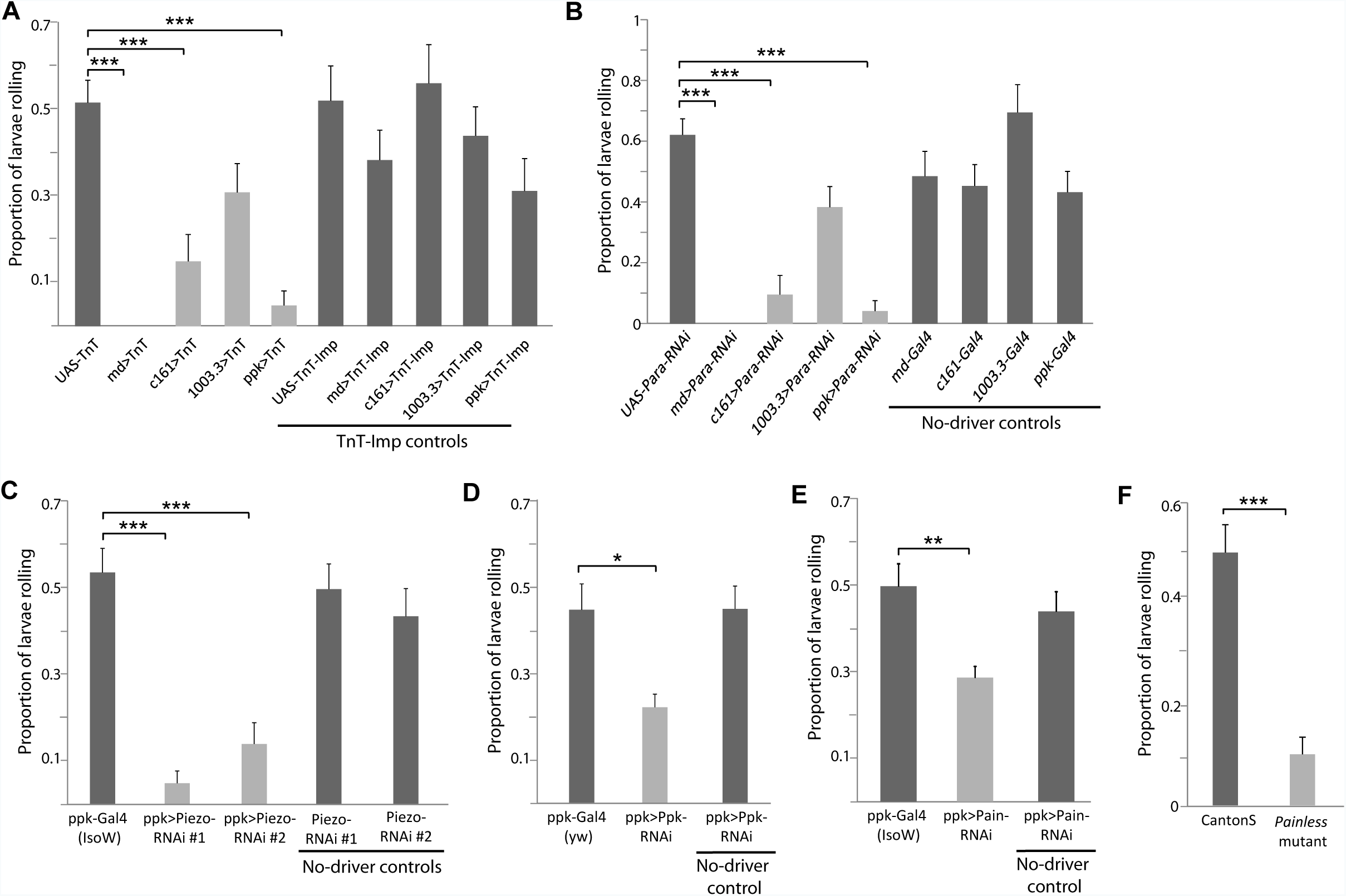
Known nociception pathway components are required for rolling in response to the gunshot stimulus. **(A-B)** Silencing different classes of multidendritic neurons with tetanus toxin light chain (TnT) but not an impotent mutant tetanus toxin (TnT-IMP) **(A)** or *para*-RNAi **(B)** reduces the larval rolling response to the gunshot. **(C-E)** RNAi against *piezo* **(C)**, *pickpocket* **(D)**, or *painless* **(E)** genes reduced the larval rolling response to gunshot. **(F)** A genetic mutation of the *painless* gene reduces the larval rolling response to BMS. (Fishers Exact Test, error bars depict standard error of the proportion *,** and *** denote significance level of Bonferonni corrected p-values, p<0.05, p<0.01, p<0.001 respectively).

Previous work has identified several genes expressed in Class IV nociceptors, including *piezo* [15], *pickpocket (ppk)* [2], and *painless (pain)* [3], to be important mediators of either mechanical (*ppk*) or polymodal nociception (*pain* and *piezo*). To determine if gunshot induced rolling relies upon these same molecular pathways, we interfered with the expression of these genes using tissue-specific RNAi or mutant larvae. We found that Class IV-specific RNAi knockdown of *piezo, pickpocket, and painless,* as well as whole-animal mutation of *painless*, significantly reduced larval sensitivity to the shot (Figure 3C-F). Together with the studies using neuronal silencing above, these findings suggest that the gunshot, like other noxious stimuli, activates nociceptive neurons embedded in the larval epidermis and relies upon dedicated nociception signaling pathways to produce rolling behavior.

We next sought to use the gunshot paradigm to conduct a screen for genes important for mechanical nociception behavior. Previous work from our laboratory has identified a set of 275 genes that are highly expressed in *Drosophila* nociceptive neurons compared to proprioceptive sensory neurons [18, 19]. This set of class IV expressed genes has been recently investigated for functions in thermal nociception[19]. We asked which of these genes play a role in the response to noxious mechanical stimuli by taking advantage of collections of UAS-driven, inverted repeat *Drosophila* lines: the GD and KK collections at the Vienna Drosophila RNAi center (VDRC), the RNAi collection at the NIG-FLY (National Institutes of Genetics, Japan), and the TRiP (Transgenic RNAi Project) collection of Harvard Medical School. We selected those lines from each collection that targeted 231 of the genes above and crossed each of these lines to the *md-GAL4;UAS-Dicer-2* and *ppk-GAL4;UAS-Dicer-2* driver stocks, thereby generating progeny with reduction in specific gene products in Class I-IV or just Class IV md neurons, respectively.

We used the gunshot method to conduct primary (Supplementary Figure 1A), secondary (Supplementary Figure 1B), and tertiary (Figure 4A) screens on the larval progeny from these crosses and quantified their rolling responses. At the time of experimentation, and during scoring of behavioral responses, the investigator was blinded to the genotypes. As an internal control for the detection of insensitive phenotypes, each day of testing included a sample of the relevant driver strain crossed to UAS-*para-RNAi.* As a control for genetic background each day of testing also included crosses to the relevant controls strains for each RNAi collection background. Here we report results of lines that that showed robust phenotypes in all three test phases of the screen. In the *md-GAL4* screen, we identified 18 lines targeting 17 genes affecting the response to the gunshot (Figure 4A, Table 1). Reduced responses were seen with RNAi against *glucose dehydrogenase* (two independent lines), *glucose transporter 1*, *dpr11*, CG14186, CG1311, *phosphomannomutase 45A, rho-like, vacuolar H+ ATPase 100kD subunit 1, lissencephaly-1,* CG10914, CG6685, CG10932, *transcription factor IIEβ, negative Cofactor 2β, mustard, and rabphilin*. RNAi against *G protein α o subunit* caused hypersensitivity.

**Table 1.**
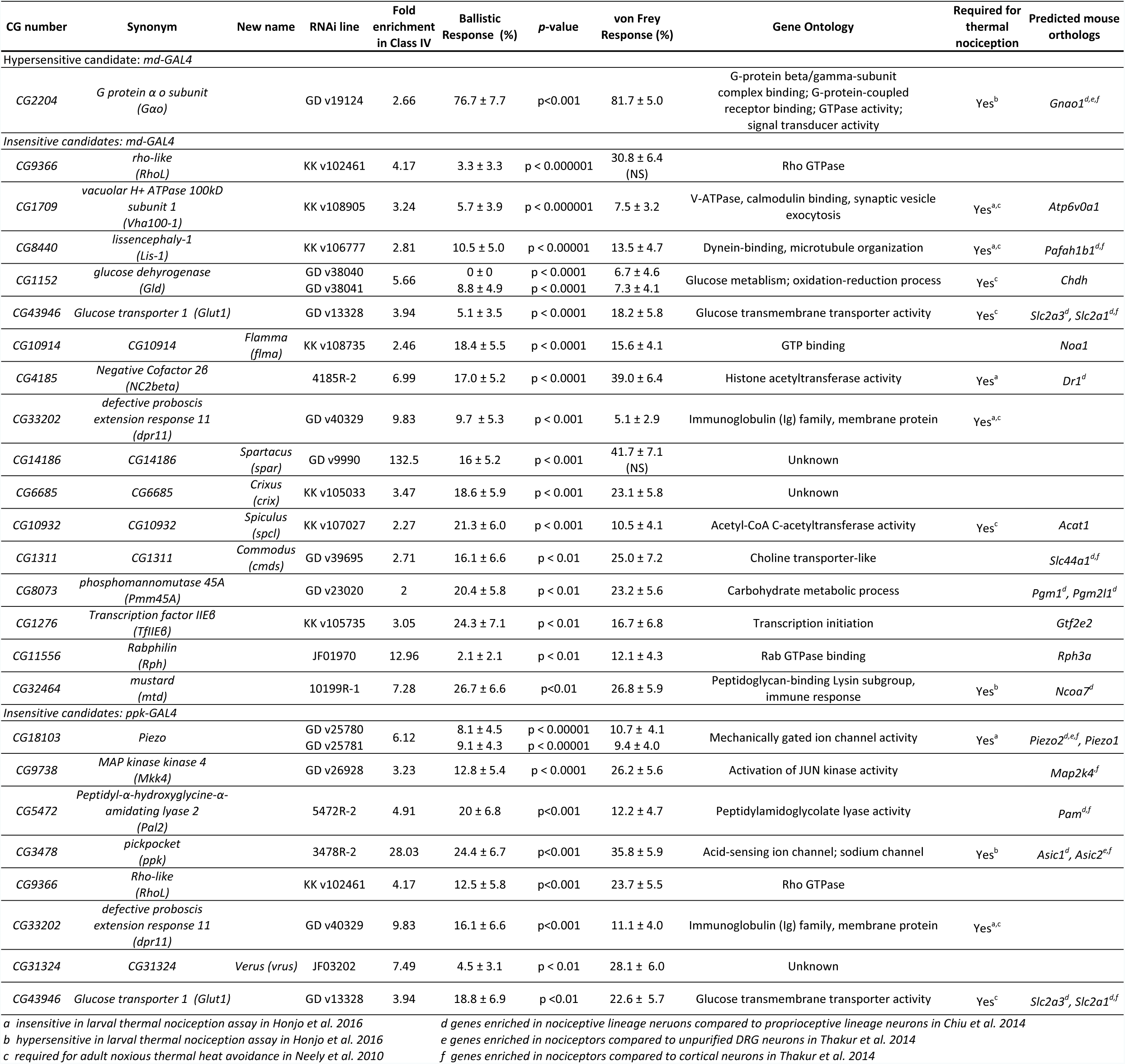
Candidate genes required for mechanical nociception responses

**Figure 4.**
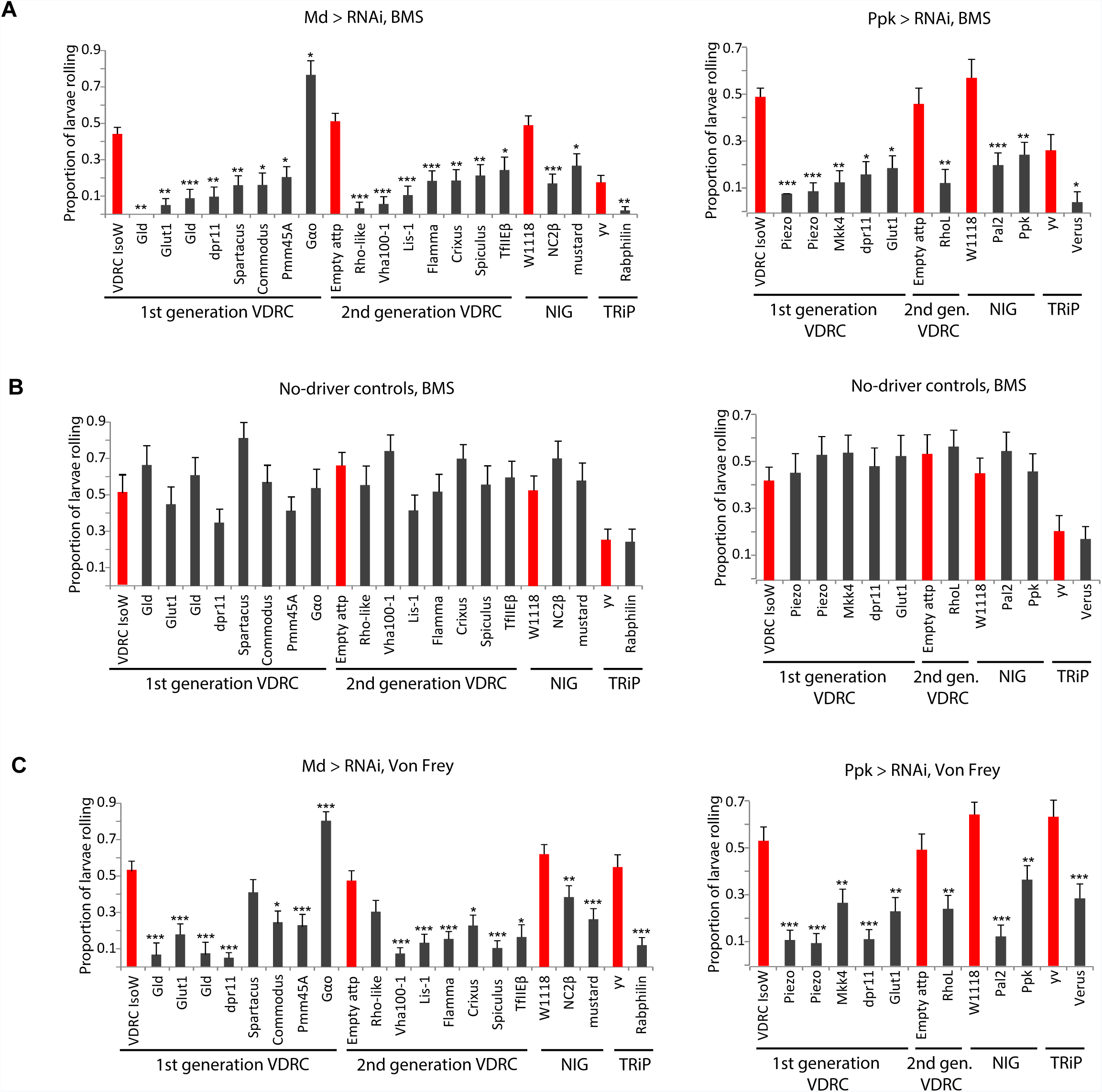
An RNAi screen using the gunshot paradigm identifies nociception genes. **(A)** Bar graphs showing the responses (total proportion from two independent testing sessions) for each RNAi line tested with the gunshot assay in the tertiary retest phase of the *md-GAL4* (*left*) and *ppk-GAL4* (*right*) screens. Control lines for each collection are shown in red. X-axis labels indicate gene names. **(B)** No-driver controls showing that Gal4 expression is necessary for the phenotypes observed in panel A for the *md-GAL4* (*left*) and *ppk-GAL4* (*right*) hits. Control lines for each collection are shown in red. X-axis labels indicate gene names. **(C)** Validation confirming that the *md-GAL4* (*left*) and *ppk-GAL4* (*right*) hits from the RNAi screen are insensitive tested in a standard manual mechanical nociception assay using von Frey fibers. Control lines for each collection are shown in red. X-axis labels indicate gene names. See also Figure S1 (Fishers Exact Test, error bars depict standard error of the proportion *,** and *** denote significance level of Bonferonni corrected p-values, p<0.05, p<0.01, p<0.001 respectively).

In the *ppk-GAL4* screen, reduced sensitivity to the gunshot was seen with 9 UAS-RNAi lines targeting 8 genes (Figure 4A, Table 1): RNAi against *piezo* (two separate lines), *MAP kinase kinase 4, dpr11, glucose transporter 1, rho-like, peptidyl-α-hydroxyglycine-α-amidating lyase 2, pickpocket, and* CG31324. All of the phenotypes from the *md-GAL4* and *ppk-GAL4* screens were driver-dependent (Figure 4B). With the exception of *rho-like* and *Spartacus,* RNAi against the candidate genes showed the expected mechanical nociception phenotypes when using the standard von Frey manual stimulation method (Figure 4C, Table 1). This latter finding is particularly important because it indicates that results with gunshot stimulation are generally comparable to results in studies using Von Frey tests.

Several of the genes that showed insensitivity to the gunshot have not been previously named. To reflect their impervious nociception phenotypes we have named these genes after famed the Roman gladiators *Spartacus (CG14186*), *Commodus (CG1311), Flamma (CG10914), Crixus(CG6685), Spiculus (CG10932),* and *Verus (CG31324)*.

Additionally, we identified a panel of genes in the *md-GAL4* and *ppk-GAL4* screens that when targeted with RNAi produced trends toward insensitive or hypersensitive phenotypes, but did not reach significance due to the stringency of the Bonferroni correction (i.e., comparisons for which p<0.05 before the Bonferroni correction)(Table S1). In the *md-GAL4* screen, RNAi against *vreteno,* CG7646, *dpr11, piezo, TNF-receptor-associated factor-like,* CG31323, CG4741, CG4398, and CG6220 produced this trend toward insensitivity, whereas RNAi against *highwire* and *snx6* produced a trend toward hypersensitivity (Table S1). In the *ppk-GAL4* arm of the screen, RNAi against *dpr11, glucose dehydrogenase, lissencephaly-1, Na+/H+ hydrogen antiporter 1, polypeptide GalNAc transferase 8, meltrin, centrosomin’s beautiful sister, CG8297, lethal (3) 03670, Flamma, and cdc42* produced an insensitivity trend, whereas RNAi against *G protein α o subunit* produced a hypersensitivity trend (Table S1).

## Discussion

We describe here a high-throughput method for the delivery of a noxious mechanical stimulus and behavioral assessment of *Drosophila* larvae. Ballistic bombardment with tungsten particles permits a simultaneous noxious stimulus to up to 50 larvae while circumventing the time-intensiveness and manual dexterity necessary for traditional methods of von Frey fiber mechanostimulation. Together, these features make the gunshot assay an ideal paradigm for the large-scale screening of larvae for behavioral defects in nociception.

The nature of the noxious stimulus by the gunshot differs substantially from that for traditional von Frey mechanostimulation methods used in *Drosophila* larvae. In the former, mechanostimulation occurs over multiple small (12µm) but discrete areas of the larval body wall due to penetration of tungsten particles in the cuticle. In the latter, mechanostimulation is confined to a single segment of the larval body wall by using a probe with a tip roughly 200 µm in diameter. However, our experiments demonstrate the reliance of the gunshot assay on the same Class IV nociceptors and a similar set of known mechanosensory genes required for response to noxious mechanical von Frey stimulation [2, 3, 11, 15]. Localized stimulation by small particles in the gunshot assay may be more representative of the ethologically-relevant stimulus during parasitoid wasp oviposition known to drive larval nociception behavior in naturalistic settings [25]. For example, the diameter of *Leptopilina* wasp ovipositors is similar (approximately 10µm when measured at the clip) to that of the 12µm particles. In addition, this natural aggressor penetrates the larval cuticle and epidermis causing a small focal area of tissue damage. It is likely that microscopic tissue damage similarly occurs with mechanical stimuli that activate mammalian nociceptors so the study of these larval sensing mechanisms may also be relevant to mammalian pain pathways.

In the gunshot paradigm, tungsten bombardment of the larval cuticle is a critical event causing larvae to roll. However, our results suggest that not every tungsten particle has the same capacity to induce rolling behavior. Indeed, even though all larvae in our paradigm are hit with multiple tungsten particles, only about half roll in response to a given bolus of tungsten (under the screening conditions used). Merely being struck with tungsten particles is thus not sufficient to induce nociceptive behavior. A possible explanation is that a certain number of particles must penetrate the cuticle to create an effective stimulus that triggers rolling. When we fired tungsten at larvae at low density, larvae that rolled had an average of at least three particles that were embedded in the cuticle (Figure 2). Non-rollers on the other hand showed only a single embedded particle on average (Figure 2). Unknown features of the ballistic stimulus, such as the size of the particle striking the larva (because individual 12-µm particles sometimes clump together to make more massive missiles), the force with which the particle reaches the larval cuticle (or, relatedly, the depth of particle penetration), or spatiotemporal pattern of stimulation by multiple particles striking the larval body wall, may each contribute to whether a given larva rolls in response. Further studies tracking particles of differing sizes, the depth of particle penetration, or the location of embedded particles relative to somata and dendrites of Class IV md neurons will shed additional light on the complexities of the stimulus evoking nociception behavior in this assay.

Our finding that particle bombardment, like von Frey fiber stimulation, evokes a nociception behavior begs the question of what actually drives nociceptor activation in response to this noxious stimulus. It is possible that these stimuli activate the nociception circuit via direct gating of mechanically-gated ion channels in Class IV md neurons [15, 26]. Alternatively, damage to the larval cuticle and epidermis could release factors that stimulate or lower the threshold for activation of multidendritic neurons [7, 8]. Given the speed of the behavioral responses any potential cell non-autonomous signals coming from these sources would need to be rapidly transmitted to Class IV dendrites. These mechanisms are neither mutually exclusive nor exhaustive, and further work is underway to clarify the neuronal transduction and activation mechanism(s) involved with this new assay.

As in our prior studies, and those in other laboratories, the activity of Class IV md neurons is necessary for triggering of nocicifensive rolling responses. Additionally, results from this study suggest that the Class II and Class III md neurons also play a role in responding to mechanically induced nociceptive behaviors (Figure 3) Therefore, we chose to pursue a dual-armed design for our behavioral screen in which we tested for mechanical nociception behavioral deficits after gene knockdown both specifically in the Class IV nociceptors as well as more broadly in all Class I-IV md sensory neurons.

We found the sets of mechanonociception genes identified as hits in these two arms of the screen to be overlapping but distinct. RNAi knockdown of three genes, including *dpr11*, *rhoL*, and *glut1,* caused significant insensitivity in both the *ppk* and *md* screens. Additionally, for several genes causing significant phenotypes in the *md-GAL4* screen (*gld, lis-1, CG10914, Vha100-1, G-α-o)*, there was a parallel (although non-significant following Bonferroni correction) trend toward hyper- or insensitivity in the *ppk-GAL4* screen. Similarly, for one gene causing significant insensitivity in the *ppk-GAL4* screen (*piezo)*, there was a trend toward insensitivity in the *md-GAL4* screen. Such concurrence of hits was not surprising, given that both arms of the screen knocked down gene products in Class IV md neurons.

We also identified hits in each screen arm that caused no significant phenotype in the other arm. While the implications of these differences require further study, they are consistent with the idea that different classes of multidendritic neurons contribute in unique ways to the detection and signaling of noxious mechanical stimuli. Genes that only showed a phenotype with knockdown by the *md-GAL4* driver may have a more widespread expression pattern and function in multiple classes of md neuron. Genes that showed a phenotype upon knockdown with *ppk-GAL4* but not with *md-GAL4* could have opposing functions in distinct classes of neuron. Alternatively, it is possible that weaker knockdown of gene expression may occur in the Class IV neurons with the *md-GAL4* driver and in some cases this may be insufficient for gene knockdown in these cells.

The set of Class IV enriched genes that we screened in this study has also been analyzed with Class IV specific knockdown and thermal nociception behavioral assays[19]. Comparisons made between these two screens allow for the identification of a set of core genes that show a general requirement in both mechanical and thermal nociception and also reveal molecules that may be uniquely required for only one or the other modality. Knockdown of *dpr-11, piezo, NC2Beta, Lis-1* and *vha-100* produced insensitive nociception in both the mechanical (this study) and thermal nociception screens [19]. Of these, reduced dendrites are seen with knockdown of *piezo, NC2Beta* and *Lis-1.* Of note, a reduced Class IV neuron dendrite phenotype is not sufficient to cause a reduced response in the gunshot assay. Knock down of genes such as *oven mitt, trivet, fire dancer* and *SECISBP2* were not found to have defective responses in the gunshot screen even though these manipulations are associated with reduced dendrites[19].

One manipulation caused hypersensitive responses in the gunshot screen (with md-GAL4 driving *G-alpha-o* RNAi). As well, *G-alpha-o* showed a trend towards hypersensitivity with the *ppk-GAL4* driver. Interestingly, *ppk-GAL4* driven RNAi against *G-alpha-o* also causes hypersensitive thermal nociception and a pronounced hyperbranched dendritic arbor in Class IV neurons. The gunshot assay identified far fewer hypersensitive phenotypes than were found with thermal nociception assays [19]. This may be a result of our use of a single testing condition that produced a behavioral nociception response in approximately 50% of animals with control genotypes. We hypothesized that this intermediate level of response would allow us to identify both insensitive and hypersensitive phenotypes in a single screen. In retrospect, it is possible that hypersensitive phenotypes would be more efficiently detected with a gunshot stimulus that is closer to threshold as this might allow for more easily observed increases in response.

As noted previously, it is important to keep in mind the caveats that are associated with the tissue specific RNAi methodologies presented here. RNAi can result in an incomplete knockdown of gene expression and phenotypes observed may be more similar to hypomorphic mutant alleles. As well, incomplete knockdown effect can also result in false negatives, which are estimated to occur in up to 40% of the UAS-RNAi strains in the major VDRC collection of strains used in our screen [27]. Thus, the lack of a phenotype in our screen cannot be used to conclusively infer a lack of function for a particular gene of interest. As well, false positives may occur, presumably due to off target effects. When the UAS-RNAi used in conjunction with *UAS-dicer-2* (as in our experiments) the effectiveness of knockdown is enhanced, and off-target effects are seen in approximately 6% of lines (when tested in the very sensitive crystalline lattice of the eye, or in the notum)[27].

Despite its significant acceleration of the behavioral stimulation process, the high-throughput paradigm described in this report still relied upon manual quantification of larval behavior after experimentation. Our preliminary efforts to automate this quantification with existing larval tracking software tools were not successful. The high density of animals in the testing arena increased throughput on the front end of screening but made following individual larvae with tracking software difficult since larvae that contacted each other were frequently lost by the tracking software. However, the continuing development of tools for the automated, high-throughput assessment of behavior in *Drosophila* is an active area of research, and future efforts to adapt existing tools or the development of new machine vision tools will greatly facilitate quantification. Such automated quantification would not only enhance the throughput of the screen but also permit the analysis of more subtle and detailed larval behaviors (e.g., latency and duration of rolling, turning, and writhing), thereby increasing our ability to detect defects in larval nociception behavior.

In summary, we have used a high-throughput mechanostimulation paradigm to identify mechanonociception genes in *Drosophila* larvae. Previously identified mechanical nociception genes are required for responses to the tungsten particle stimulus. This is consistent with the hypothesis that the behavioral responses that are triggered may be in response to forces applied by the particles. However, in addition, the stimulus penetrates the cuticle and may therefore involve signals that arise from the epidermal cells. Using this new paradigm, we have performed a genetic screen and have identified a suite of genes that are of interest for further analysis.

## Experimental Procedures

### Fly Maintenance and Stocks

Drosophila stocks were raised on standard cornmeal molasses fly food medium at 25°C and 75% humidity on a 12/12 light/dark cycle. We used the Gal4-UAS system to direct the expression of proteins or RNAi to specific neuron subtypes. The following fly stocks were used as drivers: *w*^*1118*^; *GAL4 109(2)80; UAS-dicer2* (md-GAL4, drove expression in Class I-IV md neurons), *w*^*1118*^; *ppk-GAL4; UAS-dicer2* (ppk-GAL4, drove expression in Class IV md neurons only), *w*^*1118*^; *c161-GAL4* (drove expression in Class I and II md neurons), *w*^*1118*^; *1003.3-GAL4* [28] (drove expression in Class II and III md neurons). The following effector fly stocks were used: *w; UAS-TNT* [29], *w; UAS-Tnt-IMP-V* [29], *UAS*-*para* RNAi (VDRC Transformant ID 104775), UAS-*piezo* RNAi (VDRC Transformant IDs 25780 and 25781), *UAS*-ppk RNAi (VDRC Transformant ID 108683), UAS-*painless* RNAi (VDRC Transformant ID 39478). Other fly stocks included the following: *ppk-GAL4 UAS-mCD8::GFP* and *painless*^*1*^ mutant [3]. *Drosophila* stocks used in the RNAi screen are described in Supplementary Table 1. RNAi screening stocks were provided by the Bloomington Drosophila Stock Center (TRiP collection), Vienna Drosophila RNAi Center (first and second generation collections) [55], and the National Institute of Genetics. Control stocks used for comparison with the first-generation (GD) VDRC, second-generation (KK) VDRC, TRiP, and NIG collections were VDRC isogenic w1118 (Transformant ID 60000), VDRC empty attp (VDRC Transformant ID 60100), TRiP *y*^*1*^*v*^*1*^;P{CaryP}attP2 (36303), and *w*^*1118*^, respectively.

### Gene Gun Nociception Assay

To mechanically stimulate large numbers of *Drosophila* larvae, we used a Helios gene gun system (BioRad) to deliver a small bolus of tungsten particles to larvae crawling in behavioral arenas. Briefly, 25 mg of 12-μm diameter tungsten particles (Strem Chemicals, Inc., Kehl, Germany) were suspended in ethanol and loaded into 15” segments of Tefzel tubing (BioRad, Hercules, CA) to generate the “1x” loading concentration. More dilute tungsten loading concentrations were made by decreasing the mass of tungsten particles suspended in ethanol and loaded into the 15” segments of Tefzel tubing. The suspended tungsten was spread evenly throughout the tubing segments and allowed to settle for 2-3 minutes before ethanol withdrawal. Tubing segments were then air-dried for 1-2 h and cut into individual bullets 0.5” in length for use in the gene gun system.

During testing, wandering 3^rd^ instar larvae were rinsed from the walls of vials using distilled water 5-7 days after crossing. Larvae were placed in 20-mm petri dishes containing 1 mL of distilled water. This step was sequentially performed on larvae from many genotypically distinct genetic crosses and the dishes containing the larvae of the different genotypes were arranged in order on the bench top to await testing in the assay. Larvae were never allowed to remain in the behavioral arenas for a period of greater than 3 hours prior to testing as this was the maximum time delay that we analyzed in our control experiments. Larvae from individual dishes were then positioned onto a stage 9.25” below the nozzle of the gene gun. The gene gun was loaded with the tungsten bullets described above and fired at a delivery pressure ranging from 50-110 psi, as noted in the text. Unless otherwise noted, larvae were shot using a delivery pressure of 70 psi and a tungsten loading concentration of 1x. Except where noted, larvae were shot within 1 h of rinsing from vials. Larvae in behavioral arenas were subjected to only one shot of tungsten particles.

All behavioral testing was done under red-light illumination to reduce the potential for visual cue biasing of larval behavior. The stage supporting behavioral testing arenas was illuminated using red-filtered, miniature incandescent light bulbs (40.8 W) inserted into ports drilled into the sides of the stage. A small pool of water was placed between the stage and the behavioral arena to permit efficient transmission of light from the stage to the arena. Behavioral responses to ballistic mechanical stimulation during the 30 s after stimulation were videotaped from below using a Firefly MV firewire camera (Point Grey Research, Inc, Richmond, BC, Canada) and analyzed manually offline. Responses were categorized in a binary fashion for the presence of larval NEL (rolling), which was defined as completion of a full 360° rotation. Data are reported as the proportion of larvae that exhibited NEL in response to ballistic stimulation.

### Tungsten Penetration

For experiments examining tungsten penetration into an empty behavioral arena, 35-mm petri dishes containing 1 mL of 1% agarose were shot with bullets containing 1x or 1/2x tungsten, as noted. The number of particles embedded in agarose were counted manually; data are reported as the average of 3 (for 1/2x) or 5 (for 1 x) trials. For experiments examining tungsten penetration into larvae, larvae were shot in pairs with bullets containing 1/4x tungsten. Behavioral responses were recorded, and larvae were immediately examined for particles penetrating the cuticle to minimize the likelihood of embedded particle sloughing. Penetrant particles were defined as those remaining embedded within the larval cuticle at the time of observation. To observe dendritic morphology, wandering 3^rd^ instar larvae were anesthetized with ether and mounted in glycerol. Images were taken using an Apochromat 40x (NA 1.3) oil immersion lens with a Zeiss LSM 510 Live laser scanning microscope.

### RNAi screen

For the primary phase of the screen, 6 virgin *w*^*1118*^; *md-Gal4;UAS-dicer2* or *w*^*1118*^ *ppk-Gal4; UAS-dicer2* female flies were crossed to 3 males from 461 RNAi lines and four control lines (Supplementary Table 1). Wandering third instar larval progeny from at least two independent crosses were tested using the gunshot assay on different days. Gene gun emission pressure was maintained at 70 psi during the entire screen. The proportions of larvae showing nocifensive escape locomotion in response to ballistic mechanostimulation were averaged from all days of testing, and lines with average responses greater than 1.5 standard deviations above or below the mean of all lines tested (for each of the four collections) were selected for retesting.

During the retest phase of the screen (the secondary screen), virgin *w*^*1118*^; *md-Gal4;UAS-dicer2* or *w*^*1118*^ *ppk-Gal4; UAS-dicer2* female flies were crossed to males from RNAi lines with insensitive or hypersensitive phenotypes identified during the primary screen. A minimum of 30 wandering 3^rd^ instar larval progeny from at least two independent crosses were tested on different days for each line. Those lines with Bonferroni-corrected significant insensitive or hypersensitive phenotypes were reported as hits.

Hits identified during the retest portion of the screen were run through a tripartite phase of validation. To confirm the reproducibility of the phenotypes, males from hit lines were crossed to virgin *w*^*1118*^; *md-Gal4;UAS-dicer2* or *w*^*1118*^ *ppk-Gal4; UAS-dicer2* females; wandering 3^rd^ instar larval progeny was assessed for insensitive or hypersensitivity phenotypes. To address the possibility of leaky RNAi expression, males from hit lines were crossed to virgin females from the appropriate control line (i.e., “no driver control”); wandering 3^rd^ instar larval progeny was assessed for insensitive or hypersensitive phenotypes. To test if the observed phenotypes reflect potential defects in mechanical nociception, males from candidate hit lines were crossed to *w*^*1118*^; *md-Gal4;UAS-dicer2* or *w*^*1118*^ *ppk-Gal4; UAS-dicer2* females; wandering 3^rd^ instar larval progeny was assessed for insensitive or hypersensitive phenotypes using a standard von Frey manual mechanostimulation assay as previously described [11]. In the von Frey assay, probes delivering 50 mN of maximum force were used. A minimum of 30 animals from at least two independent crosses were tested on at least two different days. Data from all testing dates were pooled, and average responses for each line were compared to controls using Bonferroni-corrected significance criteria.

### Statistics

Data were compared using a Student’s t-test (MATLAB) or Fisher’s exact probability tests (vassarstats.net), as appropriate. All statistical comparisons were done using Bonferroni-corrected significance values, with p-values less than 0.05 being considered statistically significant.

### Orthology Calls

The *Drosophila* RNAi Screening Center Integrated Ortholog Prediction Tool (DIOPT) v5.3 [30] was used to identify candidate orthologous genes from *Mus musculus*. All of the reported candidate orthologs were identified with at least two DIOPT prediction tools. Where multiple candidate orthologs were found, the highest scoring candidates were reported (Table 1).

## Acknowledgements

We are much indebted to Donald Lo for his generosity in sharing resources and experience with ballistic methods as well as for his guidance in the development of our methodology. Thanks to Dr. Ken Honjo for the assembly of the collection of RNAi lines used in the screens. Members of the Tracey lab made helpful comments on the manuscript. MGC was supported by postdoctoral fellowships from the National Institutes of Health (NIH) T32NS051156 and the Ruth K. Broad Biomedical Research Foundation. Additional support for the project was from grants to WDT (NIH R01GM086458, McKnight Foundation Technological Innovations in Neuroscience Award).

---

**Figure S1. RNAi screen using the BMS assay, related to Figure 4. (A)** Bar graphs showing the responses (average of two independent testing sessions) for each RNAi line tested with the BMS assay in the primary phase of the md (*upper*) and ppk (*lower*) screens. Lines selected for retest are highlighted in red. **(B)** Bar graphs showing the responses (total proportion from two independent testing sessions) for each RNAi line tested with the BMS assay in the retest phase of the md (*upper*) and ppk (*lower*) screens. Control lines for each collection are shown in red. X-axis labels indicate RNAi line names.

**Table S1** Summary of hits from the screen that showed a trend towards significance but which did not survive Bonferroni correction.

**Supplemental Movie 1** Representative movie showing the rolling behavioral responses of a wild type control strain to the ballistic stimulus. The timing of the shot is indicated approximately 2 seconds into the video.

## Literature Cited

1. Hwang, R.Y. (2009). Circuitry and Genes of Larval Nociception in Drosophila Melanogaster. PhD Dissertation, Duke University http://hdl.handle.net/10161/1348.

2. Hwang, R.Y., Zhong, L., Xu, Y., Johnson, T., Zhang, F., Deisseroth, K., and Tracey, W.D. (2007). Nociceptive neurons protect Drosophila larvae from parasitoid wasps. Curr Biol 17, 2105–2116.

3. Tracey, W.D., Wilson, R.L., Laurent, G., and Benzer, S. (2003). painless, a Drosophila Gene Essential for Nociception. Cell 113, 261–273.

4. Story, G.M., Peier, A.M., Reeve, A.J., Eid, S.R., Mosbacher, J., Hricik, T.R., Earley, T.J., Hergarden, A.C., Andersson, D.A., Hwang, S.W., et al. (2003). ANKTM1, a TRP-like channel expressed in nociceptive neurons, is activated by cold temperatures. Cell 112, 819–829.

5. Zhong, L., Bellemer, A., Yan, H., Honjo, K., Robertson, J., Hwang, R.Y., Pitt, G.S., and Tracey, W.D. (2012). Thermosensory and non-thermosensory isoforms of Drosophila melanogaster TRPA1 reveal heat sensor domains of a thermoTRP channel. Cell Reports 1, 43–55.

6. Caterina, M.J., Schumacher, M.A., Tominaga, M., Rosen, T.A., Levine, J.D., and Julius, D. (1997). The capsaicin receptor: a heat-activated ion channel in the pain pathway. Nature 389, 816–824.

7. Babcock, D.T., Landry, C., and Galko, M.J. (2009). Cytokine signaling mediates UV-induced nociceptive sensitization in Drosophila larvae. Curr Biol 19, 799–806.

8. Babcock, D.T., Shi, S., Jo, J., Shaw, M., Gutstein, H.B., and Galko, M.J. (2011). Hedgehog Signaling Regulates Nociceptive Sensitization. Curr Biol.

9. Im, S.H., Takle, K., Jo, J., Babcock, D.T., Ma, Z., Xiang, Y., and Galko, M.J. (2015). Tachykinin acts upstream of autocrine Hedgehog signaling during nociceptive sensitization in Drosophila. eLife 4, e10735.

10. Neely, G.G., Keene, A.C., Duchek, P., Chang, E.C., Wang, Q.P., Aksoy, Y.A., Rosenzweig, M., Costigan, M., Woolf, C.J., Garrity, P.A., et al. (2011). TrpA1 Regulates Thermal Nociception in Drosophila. PLoS One 6, e24343.

11. Zhong, L., Hwang, R.Y., and Tracey, W.D. (2010). Pickpocket is a DEG/ENaC protein required for mechanical nociception in Drosophila larvae. Curr Biol 20, 429–434.

12. Neely, G.G., Hess, A., Costigan, M., Keene, A.C., Goulas, S., Langeslag, M., Griffin, R.S., Belfer, I., Dai, F., Smith, S.B., et al. (2010). A Genome-wide Drosophila Screen for Heat Nociception Identifies alpha2delta3 as an Evolutionarily Conserved Pain Gene. Cell 143, 628–638.

13. Ohyama, T., Schneider-Mizell, C.M., Fetter, R.D., Aleman, J.V., Franconville, R., Rivera-Alba, M., Mensh, B.D., Branson, K.M., Simpson, J.H., Truman, J.W., et al. (2015). A multilevel multimodal circuit enhances action selection in Drosophila. Nature 520, 633–639.

14. Hwang, R.Y., Stearns, N.A., and Tracey, W.D. (2012). The Ankyrin Repeat Domain of the TRPA Protein Painless Is Important for Thermal Nociception but Not Mechanical Nociception. PLoS One 7, e30090.

15. Kim, S.E., Coste, B., Chadha, A., Cook, B., and Patapoutian, A. (2012). The role of Drosophila Piezo in mechanical nociception. Nature 483, 209–212.

16. Gorczyca, D.A., Younger, S., Meltzer, S., Kim, S.E., Cheng, L., Song, W., Lee, H.Y., Jan, L.Y., and Jan, Y.N. (2014). Identification of Ppk26, a DEG/ENaC Channel Functioning with Ppk1 in a Mutually Dependent Manner to Guide Locomotion Behavior in Drosophila. Cell Reports 9, 1446–1458.

17. Guo, Y., Wang, Y., Wang, Q., and Wang, Z. (2014). The role of PPK26 in Drosophila larval mechanical nociception. Cell Reports 9, 1183–1190.

18. Mauthner, S.E., Hwang, R.Y., Lewis, A.H., Xiao, Q., Tsubouchi, A., Wang, Y., Honjo, K., Skene, J.H.P., Grandl, J., and Tracey, W.D. (2014). Balboa Binds to Pickpocket In Vivo and Is Required for Mechanical Nociception in Drosophila Larvae. Curr Biol 24, 2920–2925.

19. Honjo, K., Mauthner, S.E., Wang, Y., Skene, J.H., and Tracey, W.D., Jr. (2016). Nociceptor-Enriched Genes Required for Normal Thermal Nociception. Cell Reports 16, 295–303.

20. Caldwell, J.C., and Tracey, W.D., Jr. (2010). Alternatives to mammalian pain models 2: using Drosophila to identify novel genes involved in nociception. Methods Mol Biol 617, 19–29.

21. Rizki, T.M., Rizki, R.M., and Carton, Y. (1990). Leptopilina-Heterotoma and L-Boulardi - Strategies to Avoid Cellular Defense Responses of Drosophila-Melanogaster. Exp Parasitol 70, 466–475.

22. Rizki, R., and Rizki, T. (1984). Selective destruction of a host blood cell type by a parasitoid wasp. PNAS 81, 6154–6158.

23. Rizki, T.M., Rizki, R.M., and Grell, E.H. (1980). A mutant affecting the crystal cells in Drosophila melanogaster. Roux’s Arch Dev Biol 188, 91–99.

24. Rizki, M.T., and Rizki, R.M. (1959). Functional significance of the crystal cells in the larva of Drosophila melanogaster. J Biophys Biochem Cytol 5, 235–240.

25. Robertson, J.L., Tsubouchi, A., and Tracey, W.D. (2013). Larval defense against attack from parasitoid wasps requires nociceptive neurons. PLoS One 8, e78704.

26. Coste, B., Xiao, B., Santos, J.S., Syeda, R., Grandl, J., Spencer, K.S., Kim, S.E., Schmidt, M., Mathur, J., Dubin, A.E., et al. (2012). Piezo proteins are pore-forming subunits of mechanically activated channels. Nature.

27. Dietzl, G., Chen, D., Schnorrer, F., Su, K.C., Barinova, Y., Fellner, M., Gasser, B., Kinsey, K., Oppel, S., Scheiblauer, S., et al. (2007). A genome-wide transgenic RNAi library for conditional gene inactivation in Drosophila. Nature 448, 151–156.

28. Hughes, C.L., and Thomas, J.B. (2007). A sensory feedback circuit coordinates muscle activity in Drosophila. Mol Cell Neurosci 35, 383–396.

29. Sweeney, S.T., Broadie, K., Keane, J., Niemann, H., and O’Kane, C.J. (1995). Targeted expression of tetanus toxin light chain in Drosophila specifically eliminates synaptic transmission and causes behavioral defects. Neuron 14, 341–351.

30. Hu, Y., Flockhart, I., Vinayagam, A., Bergwitz, C., Berger, B., Perrimon, N., and Mohr, S.E. (2011). An integrative approach to ortholog prediction for disease-focused and other functional studies. BMC bioinformatics 12, 357.

